# Inferring ecological selection from multidimensional community trait distributions along environmental gradients

**DOI:** 10.1101/2023.10.10.561738

**Authors:** Elina Kaarlejärvi, Malcolm Itter, Tiina Tonteri, Leena Hamberg, Maija Salemaa, Päivi Merilä, Jarno Vanhatalo, Anna-Liisa Laine

**Author notes:** Co-First Authors, equal contribution (Elina Kaarlejärvi, Malcolm Itter).

## Abstract

Understanding the drivers of community assembly is critical for predicting the future of biodiversity and ecosystem services. While trait-based frameworks are often used to this end, correlations among traits and the assumption of unimodal trait-abundance distributions confound detection of the underlying processes. To overcome these caveats, here we quantify multidimensional trait distributions of communities (community trait niches), which we use to identify ecological selection types shaping communities along environmental gradients. We find that directional, stabilizing, and divergent selection all modify community trait niches in over 3600 boreal forest understory plant communities, and selection on a particular trait may change from one type to another over time. Our results provide novel and rare empirical evidence from a natural system for divergent selection. The results also show that while higher trait diversity (measured as trait niche volume) is generally associated with higher species richness, high resource availability may enable tighter niche packing. Jointly our approach provides a framework for identifying key traits under selection and facilitates the detection of processes underlying community dynamics.

## Introduction

Under global environmental change, it is increasingly important to understand how natural communities and their provision of ecosystem services are changing. Communities are composed of individuals with different phenotypic characteristics influencing their performance, *i*.*e*. their functional traits (McGill et al. 2006, Violle et al. 2007). The abiotic environment and biotic interactions select for individuals conditional on their traits. When selection favours individuals with a certain trait value, regardless of species, we can expect a subsequent increase of this trait value in the community (Shipley 2010, Vellend 2010, Shipley et al. 2016). Communities and their trait composition are thus aggregate properties stemming from individuals with different trait combinations responding to varying selection pressures. As individuals express multiple traits, selection on one trait inevitably leads to concomitant shifts in other traits, and may thus alter the correlation among traits at the community level (Laughlin 2014, Kraft et al. 2015b, Clark 2016, Dwyer and Laughlin 2017). Nevertheless, the community assembly literature is dominated by studies, which focus on single traits and frequently find evidence of abiotic rather than biotic processes structuring communities (Götzenberger et al. 2012, Münkemüller et al. 2020). Moreover, the commonly used filtering framework, which compares observed variation in single traits to variation produced by random assembly, is often unable to disentangle abiotic and biotic processes, thus providing limited transferable knowledge on how communities are structured (Münkemüller et al. 2020). Identification of ecological selection types offers a more general view on assembly of communities (Vellend 2010, 2016). Recent studies have explored how selection impacts the distributions of single traits in communities (Vellend 2016, Rolhauser and Pucheta 2017, Loranger et al. 2018, Gross et al. 2021, DeMalach et al. 2022). Here, we expand this approach to account for multiple correlated traits – a necessity for more reliable inference on assembly processes (Kraft et al. 2015b, Clark 2016, Wüest et al. 2018, Pistón et al. 2019) – and identify different selection types driving community composition along environmental gradients.

To date few studies have predicted multidimensional trait diversity patterns (Carmona et al. 2016, Clark 2016, Blonder 2018), as multi-trait approaches have been limited by challenges with modelling trait correlations and comparing multidimensional trait volumes (Carmona et al. 2016, Blonder 2018, Lu et al. 2021). Recent joint-trait methods include ordinations (Mouillot et al. 2021), the dynamic adaptive landscape framework (Laughlin and Messier 2015), multivariate kernel density estimation (Blonder et al. 2014) and boosted regression trees (Elith et al. 2008). The Trait Probability Density framework by Carmona *et al*. (2016) introduces a probabilistic way to quantify multidimensional trait hypervolumes. We transform this conceptual probabilistic framework into a predictive one by first modelling species relative abundances along environmental gradients and then translating them to multidimensional trait distributions. Our model fully accounts for trait correlations (or trait syndromes) imposed by species phenotypes, because we model species’ – not single or separate traits’ – responses to environment. For this we use a multinomial Bayesian joint species distribution model (JSDM) (Itter et al. 2023), which accurately predicts observed species relative abundances (Itter et al. 2023). We then use posterior model predictions to construct probabilistic multivariate trait distributions following a modified version of the predictive trait model introduced by Clark (Clark 2016). As our JSDM does not account for ecological interactions between species in a causal or mechanistic way, as is true for any JSDM (Blanchet et al. 2020, Poggiato et al. 2021), we do not use this model to separate effects of abiotic vs. biotic selection. Instead, by investigating patterns emerging in the predicted multidimensional trait distributions of communities (hereafter community trait niches) along environmental gradients, we demonstrate how this framework can be used to infer ecological selection types and detect potential low-level processes driving community dynamics, contributing to the recent development in trait-based ecology (Webb et al. 2010, Laughlin 2014, Dwyer and Laughlin 2017, Gross et al. 2021).

Here, we focus on species-poor boreal forest understory communities to determine the types of ecological selection and potential mechanisms modifying community diversity and composition. By examining changes in community trait niches along modelled environmental gradients, we address three objectives: First, we assess whether community trait niche volume is associated with species richness. Following reasoning of limiting similarity hypothesis stating that coexisting species cannot occupy the same niches (Macarthur and Levins 1967), we expect that higher species richness will require a larger niche volume (Fig. 1a). Second, we investigate which traits contribute most to overall change in the trait niche volume (increase or decrease trait variation) along environmental gradients, *i*.*e*. identify the traits under selection (Fig. 1b). Since nitrogen limitation intensifies along the successional gradient in boreal forests (Vitousek and Reiners 1975, Merilä et al. 2002, Högberg et al. 2017), we expect nutrient acquisition related traits such as plant foliar carbon to nitrogen ratio (C:N) and leaf dry matter content (LDMC) to be under selection along the successional gradient. Third, we assess how bivariate and marginal trait distributions (*i*.*e*., cross-sections of the multidimensional niches) change along modelled environmental gradients to detect signs of three types of ecological selection (Vellend 2016) (Fig. 1c). Directional selection leads to a shift in the most common, and thereby presumably optimal (Violle et al. 2007, Enquist et al. 2015), trait values in the community (Fig. 1d), driven e.g. by environmental filtering (Keddy 1992, Kraft et al. 2015a) or competitive exclusion (Mayfield and Levine 2010) (Table 1). Stabilizing selection reduces trait variation and results in trait convergence evidenced by a narrower range of traits (Vellend 2010, 2016, Rolhauser and Pucheta 2017, Loranger et al. 2018, DeMalach et al. 2022) (Fig. 1e). It can be driven by two opposite environmental filters acting on the same trait (Kraft et al. 2015a, Rolhauser and Pucheta 2017), by competitive exclusion (Mayfield and Levine 2010), or by positively frequency dependent mechanisms (Vellend 2016), which benefit species with similar response traits. For example during succession some plants may influence soil microbial composition so that it facilitates other plant species with similar traits (environmental requirements) to colonize and gain abundance (van der Putten et al. 2013, Fukami 2015). Stabilizing selection may also increase trait correlations without a corresponding change in single trait means or variances (Lande and Arnold 1983, Arnold et al. 2001) (Fig. 1f). This stronger trait correlation may arise from an environmental filter such as aridity (Dwyer and Laughlin 2017), or biotic interactions such as predation (Mikolajewski et al. 2010, Tanaka 2012). Finally, divergent selection leads to an increase in the number of modes (‘peaks’) in trait distributions, or to more even trait distributions (Kingsolver and Pfennig 2007, Vellend 2016, Gross et al. 2021) (Fig. 1g). Divergent selection may be generated by limiting similarity (Macarthur and Levins 1967, Weiher and Keddy 1995, Meszéna et al. 2006) or negative species interactions, such as resource competition via niche differences (Levine and HilleRisLambers 2009, Mayfield and Levine 2010) or apparent competition for example via shared pathogens or mutualists (Allan et al. 2010, Devaux and Lande 2010, Siefert et al. 2018) (Table 1). We note that bimodality or a uniform trait distribution can also emerge from a sudden shift in directional selection, if a new trait optimum has appeared but the individuals representing the past optimum are still present.

**Table 1.** Three types of ecological selection, their consequences on trait distributions, examples of underlying low-level mechanisms, classification of the mechanisms into abiotic and biotic filters and references for mechanisms.

| Type of selection | Pattern in trait distribution | Examples of low-level mechanisms | Filter | References |
| --- | --- | --- | --- | --- |
| Directional | Shift in mean | Environmental filter<br>Competitive exclusion | Abiotic<br>Biotic | (Keddy 1992, Kraft et al. 2015a)<br>(Mayfield and Levine 2010) |
| Stabilizing | Reduced variation in single trait | Competitive exclusion<br>Two opposite env. filters<br>Positively frequency dependent mechanisms:<br>Priority effects<br>Facilitation | Biotic<br>Abiotic<br>Biotic<br>Biotic<br>Biotic | (Mayfield and Levine 2010)<br>(Kraft et al. 2015a, Rolhauser and Pucheta 2017)<br>(Vellend 2010)<br>(Fukami 2015)<br>(van der Putten et al. 2013) |
| Stabilizing | Stronger correlation among traits | Trade-offs between traits caused by<br>Environmental filter<br>Species interactions | Abiotic<br>Biotic | (Kraft et al. 2015a, Dwyer and Laughlin 2017)<br>(Mikolajewski et al. 2010, Tanaka 2012) |
| Divergent | >1 modes | Negatively frequency dependent mechanisms:<br>Resource competition<br>Apparent competition<br>Limiting similarity<br>Facilitation | Biotic<br>Biotic<br>Biotic<br>Biotic<br>Biotic | (Vellend 2016)<br>(Levine and HilleRisLambers 2009, Mayfield and Levine 2010)<br>(Allan et al. 2010, Devaux and Lande 2010, Siefert et al. 2018)<br>(Macarthur and Levins 1967, Weiher and Keddy 1995, Meszéna et al. 2006)<br>(Spasojevic and Suding 2012) |

**Figure 1.**
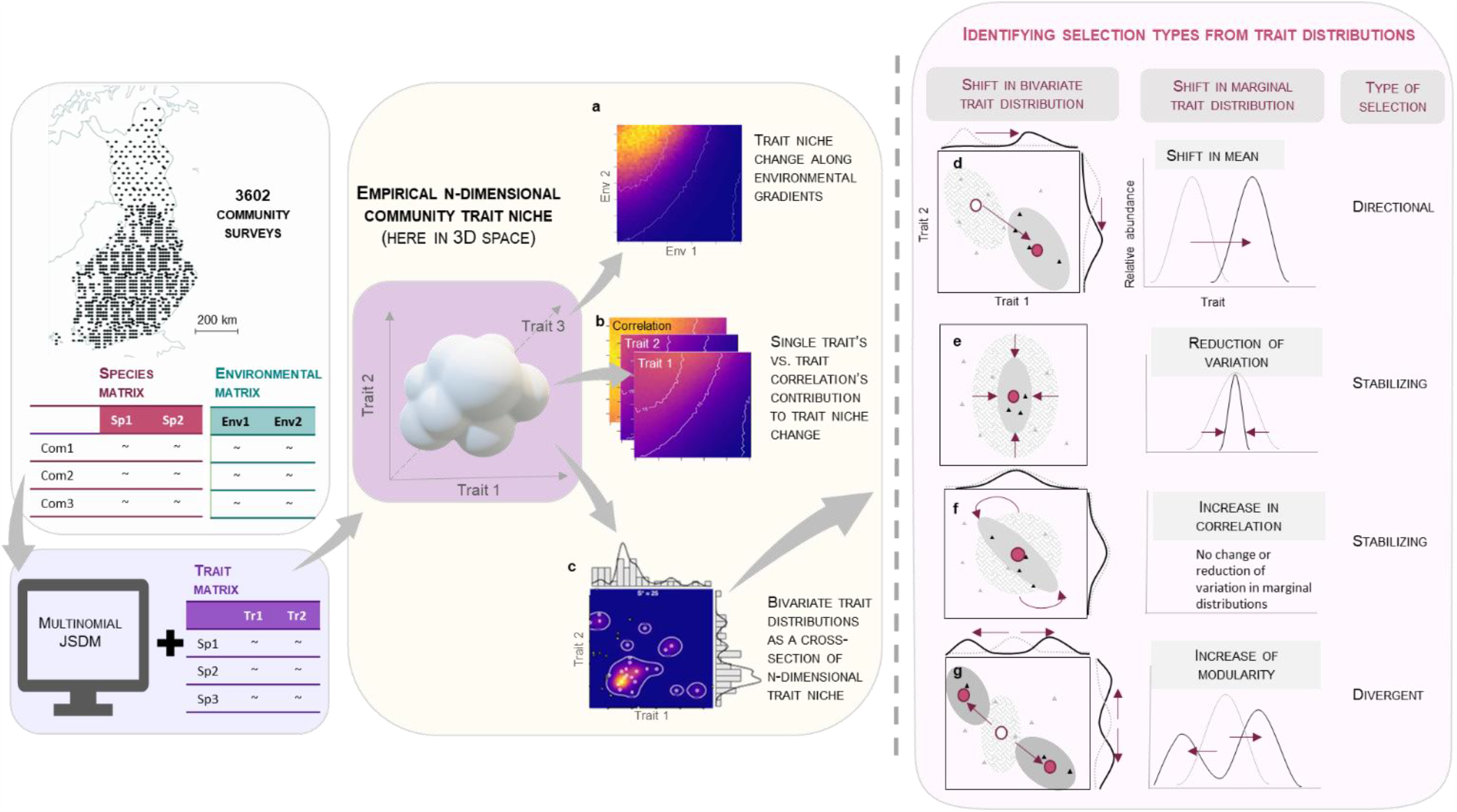
Schematic representation of the workflow and hypotheses. Empirical *n*-dimensional community trait niches were built by transforming species probability estimates (from JSDM) into multivariate trait probability distributions. Using these community trait niches, we: (**a**) investigated the relationship between trait niche volume and species richness along the modelled environmental gradients (colour represents the relative change in volume, contours illustrate richness): (**b**) partitioned trait niche change among individual traits and their correlation along the environmental gradients; and (**c**) used cross-sections of trait niches (*i*.*e*., bivariate trait distributions) to identify different selection types and potential underlying mechanisms (Table 1). Identification of ecological selection types based on shifts in mean, variance, correlation and number of modes are illustrated both for the marginal distribution of a single trait and for bivariate trait distributions (**d-g**, see Table 1). The arrows in the subpanels illustrate one possible direction. The weaved and grey ellipsoids represent the trait niche space occupied by communities in the hypothetical environments A and B, respectively. The red point represents the centroid of the space and triangles are species-specific trait values. Note that species have fixed locations in the trait space, defined by their mean trait values (*i*.*e*., trait syndrome; intraspecific trait variation ignored here). Black triangles illustrate species present in environment B; grey triangles illustrate species in the species pool, but absent from the environment B. The grey and black distributions in the margins of panels **d** and **g** a represent the shifts in single trait distributions when moving from environment A to B.

We used a JSDM to predict how growing season temperature, habitat fertility and forest density jointly influence the relative abundance of 39 vascular plant species (model specifications in Appendix S1: Section S1, species list in Appendix S1: Table S1). The data covers over 3500 understory communities across a 1200-km latitudinal gradient in Finland, sampled in 1985, 1995 and 2006. The predicted relative abundances of species were mapped to empirical trait distributions using ten traits: height (related to dispersal and ability to compete for light), specific leaf area (SLA, related to light and nutrient capture), leaf carbon to nitrogen ratio (C:N; describing relative nitrogen acquisition), leaf dry matter content (LDMC, related to nutrient acquisition), leaf phosphorus (P, related to phosphorus acquisition), leaf N (related to nitrogen acquisition), leaf N:P ratio (describing relative nutrient requirements), mycorrhizal colonization rate (describing association with soil fungi, measured as a proportion of fine root colonized by mycorrhizal fungi), clonality (binary; describing whether species is able to spread clonally) and dispersal mode (binary; describing whether species is wind dispersed). Finally, we used the four continuous traits (SLA, LDMC, leaf C:N, leaf P) to build empirical 4-dimensional trait niches at the community-level and investigated how these community trait niches changed along the modelled environmental gradients. These four traits were best predicted by the model (Appendix S1, Fig. S1).

## Materials and Methods

### Understory community and forest stand data

Understory communities were surveyed by National Resources Institute Finland (Luke) in 1985-1986, 1995 and 2006 on a systematic network of 1,700 sites established on mineral-soil in forested land. The systematic sampling network is part of the 8th Finnish National Forest Inventory (Reinikainen et al. 2000) and consists of clusters, which were located 16 km from each other in southern Finland, and 24 and 32 km apart in northern Finland along east–west and north–south axes, respectively. Each cluster includes four linearly located sampling sites 400 m apart from each other in southern Finland and three sampling sites 600 m apart from each other in northern Finland. Data includes 1495 sites in 1985-1986, 1673 sites in 1995 and 439 sites in 2006 (3602 unique inventory site-by-year combinations in total). The survey in 2006 was part of the BioSoil project carried out under the Forest Focus scheme, which is a subset of the pan-European International Co-operative Programme on Assessment and Monitoring of Air Pollution Effects on Forests (UN-ECE ICP Forests) monitoring site network (Level I) (Lorenz and Fischer 2013). Then the spatial extent of this sampling was comparable to previous surveys covering the whole country (Fig. 1), but at maximum 1 site per cluster was surveyed.

In all three surveys, within each site, vascular plant cover (0.1–100%) was visually estimated for each species on four permanent square-shaped sampling quadrats of 2 m^2^, located 5 m apart from each other. Here we modelled data at site-level and used the average of species cover across the four sampled quadrats as an estimate of species abundance at each site with percent cover values treated as species-level counts. In total 380 vascular plant species were observed in the surveys. To ensure meaningful estimation of species-specific model parameters, we focused on 39 species, which occurred in at least five percent of sites in each inventory year (a common practice in high-dimensional JSDM settings (Clark et al. 2017)).

At each site, forest density was estimated as the basal area of the trees (Tomppo et al. 2011). Shrub cover was measured as the projected percent cover of shrubs and 0.5-1.5 m tall trees within an 9.8 m radius (11.3 m in 2006) circular plot centered on the permanent vegetation survey sites. Site fertility was determined in the field using six classes based on vegetation (Cajander 1949, Pohjanmies et al. 2021). Here, we re-classified these into two groups representing ‘high’ and ‘low’ fertility. The growing degree day variable represents the site-level average annual sum of daily mean temperatures exceeding +5°C over the preceding decade calculated based on 10 km^2^-resolution interpolated daily temperature values modeled by the Finnish Meteorological Institute (Venäläinen et al. 2005).

### Trait data

Species-specific values for height, SLA, LDMC were obtained as median from the TRY database (Kattge et al. 2011); leaf C:N primarily from own measurements and secondarily as median from TRY; leaf N, P and N:P primarily from own measurements and secondarily as median from NP database (Tian et al. 2019); clonality from CLO-PLA database (Klimešová et al. 2017); mycorrhizal colonization as median from GRooT database (Guerrero-Ramírez et al. 2021); and wind pollination syndrome from TRY (pollination syndrome re-classified here to binary: wind vs. other). Own nutrient measurements were based on a pooled sample of leaves (ca. 2g) collected from multiple individuals across several locations in Southern and Eastern Finland. The samples were dried at +60°C for 72h, milled and analyzed for C and N using flow-injection analyses and for P spectrophotometrically after burning at +550°C and extracting with hydrochloric acid.

### Modelling approach

Functional trait prediction under varying environmental conditions occurs as a two-stage process. In the first stage, a multinomial Bayesian joint species distribution model (JSDM) is fit to species relative ab *Functional trait prediction* undances. In the second stage, the parameterized JSDM is used to predict species’ probabilities of success under specified environmental conditions, which are then mapped to traits through a linear transformation. Full description of the multinomial JSDM for predicting species relative abundances along growing season temperature, habitat fertility and forest density gradients can be found in Appendix S1: Section S1. The model applies a multinomial response function, which estimates each species probability of success under local environmental conditions relative to other species in the community. This means that we model community structure, not only individual species responses. Each species’ relative abundance is influenced by other species’ responses to the modelled environmental variables as well as to random (*i*.*e*., unmeasured environmental) variation between sites. The model assumes finite space and resources: if the probability of one species increases in a community, the total probability of success of all other species must decrease. While this assumption reflects antagonistic interactions among species within the same trophic level – due to shared and limited resources, e.g., nutrients, light and growing space, our model is not mechanistic and does not explicitly account for biotic interactions. Rather, the multinomial response function accounts for the non-independence of the relative abundances of species within the modelled communities. This statistical dependence is commonly modelled through a residual random effect term in existing JSDMs (Clark et al. 2017, Ovaskainen et al. 2017, see e.g. Poggiato et al. 2021). We explored the inclusion of such residual dependence here but found the multinomial response function alone led to better predictions of observed communities (Itter et al. 2023). Our modelling approach focuses thus on ecological selection (Vellend 2016) by accounting for several key environmental selection factors and dependence among species within a community.

### Functional trait prediction

The fitted multinomial JSDM was applied to predict the expected probability of success of each species under specified environmental conditions. Specifically, posterior predictions of the linear predictor for each species (*j* = 1, …, *m*) are generated as, 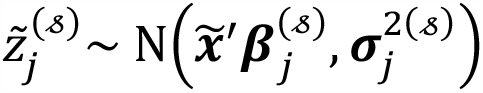, where 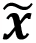 specifies the predictive environmental conditions, ***β***_*j*_ is a vector of species-specific responses to environmental conditions, 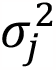 is a species-specific variance term, and 𝓈 indexes the posterior sample (𝓈 = 1, …, 𝒮). The expected probability of success of a given species under the specified environmental conditions is generated conditional on 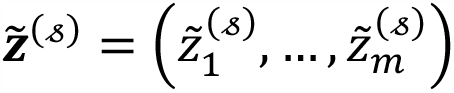 as, 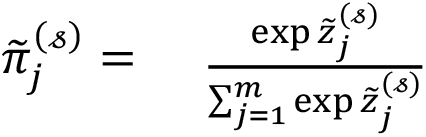.

Mapping predicted species probabilities to underlying functional traits is achieved through a linear transformation. The expectation and variance of a local community under the multinomial model are given by:

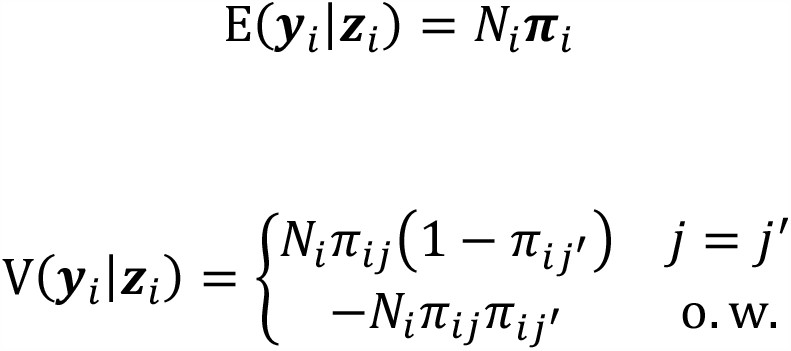

where E(⋅) and V(⋅) denote expectation and variance, respectively, and *N*_*i*_ indicates the total abundance for *i*th local community. Setting *N*_*i*_ = 1, the community weighted trait mean (CWM) and variance (CWV) are calculated as, CWM_i_ = ***T*′**E(***y***_*i*_|***z***_*i*_) = ***T*′*π***_*i*_ and CWV_i_ = ***T*′**V(***y***_*i*_|***z***_*i*_)***T***, where ***T*** is an (*m* × 𝓁)-dimensional trait matrix defined for *m* species and 𝓁 functional traits. Inserting posterior predictions of species probabilities 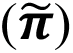 into the formulas for the community weighted trait mean and variance allows for the prediction of community-level trait distribution parameters under specified environmental conditions: 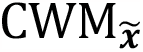 and 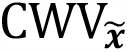. Inherent trait syndromes expressed as non-zero covariance among traits (non-independence of traits) is captured through the modeled covariance among species possessing specific trait values as quantified by the covariance matrix ***T*′**V(***y***_*i*_|***z***_*i*_)***T***.

### Baseline community

For reference, we define a baseline community in which all *m* species have equal probability of occurrence: 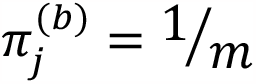 where (*b*) indicates baseline. This artificial community without any selection (Shipley 2010) provides a constant reference point to which we measure departures of predicted communities using e.g. CWM, CWV and trait hypervolume size and dimensions (see below). The baseline community is not to be confused with a null community, which requires decisions on randomization technique to sample species from the regional pool.

### Functional trait hypervolumes

We utilize the methodology presented in Lu et al., (2021) to estimate predicted trait hypervolumes (here called community trait niches) and contribution of each trait and their correlation to changes in hypervolume along environmental gradients. Specifically, functional trait hypervolume is calculated as the determinant of the 𝓁-dimensional community weighted trait covariance matrix: |CWV|. Conditional on posterior predictions of the community weighted trait covariance matrix (see Functional trait prediction), we generate predictions of functional trait hypervolumes under specified environmental conditions. For reference, predicted functional trait hypervolumes are presented relative to the hypervolume of the baseline community, 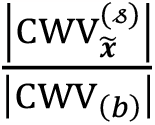, and report the posterior median ratio. Note, ratio values larger than one indicate greater hypervolume under specified environmental conditions than in the baseline community, while values less than one indicate reduced hypervolume relative to the baseline community. Given that the community weighted trait covariance matrix is a positive definite matrix, its determinant is always positive thereby ensuring the hypervolume ratio is also positive.

The determinant of the community weighted trait covariance matrix is given by,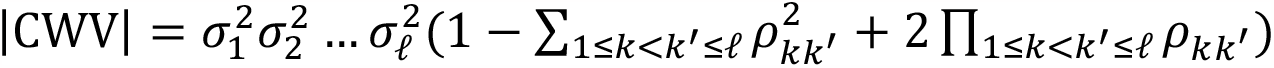, where 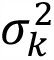 is the variance of trait *k* and *ρ*_*kk*′_ is the correlation between traits *k* and *k*′ (*k* = 1, …, 𝓁). As described in Lu et al., (2021), we can use the determinant formula to partition changes in the trait hypervolume to multiplicative terms corresponding to individual trait variances (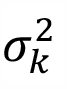 for *k* = 1, …, 𝓁) and their correlation. The correlation component is defined by all terms within parentheses, 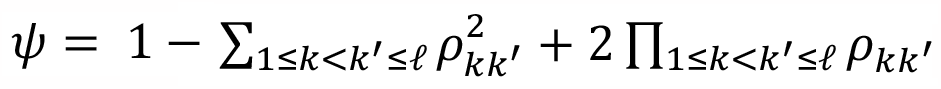, and reaches a maximum value of 1 when all traits are orthogonal such that their correlation is zero. The greater the correlation among traits the smaller the value of *ψ*. We calculate relative change in trait-level variance as, 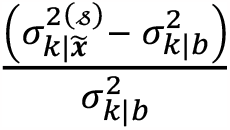 where 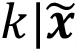 and *k*|*b* indicate the *k*th trait for the specified environment and baseline community, respectively. Similarly, we calculate the relative change in the correlation component as: 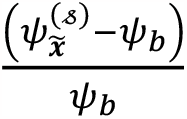. For variance terms, values less than zero indicate a decrease in trait-level variance, while positive values indicate an increase in trait-level variance relative to the baseline community. For the correlation component, values less than zero indicate a decrease in trait independence (*i*.*e*., an increase in trait dependence) potentially shrinking the trait hypervolume. Values greater than zero indicate an increase in trait independence (decrease in dependence). We report the posterior median change in individual trait variation and the correlation component.

### Empirical trait distributions

Predicted species probabilities under specified environmental conditions 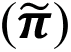 are used to construct empirical trait distributions. Note that these probabilities provide the relative weights of each species’ mean trait value within a predicted local community (see Functional trait prediction). Empirical trait distributions are constructed by specifying a total community abundance (*N*) and sampling the mean trait values of constituent species with frequencies equal to predicted species probabilities. All empirical trait distributions utilize the median of the posterior predictive species probability vector under specified environmental conditions. We then compute univariate and bivariate Gaussian kernel density estimates of the sampled functional trait values using the minimum and maximum trait values as the density bounds (i.e., density is equal to zero beyond these values). In bivariate plots, ellipsoids are generated for select quantiles applying linear interpolation to the cumulative sum of the empirical trait density. Baseline empirical trait distributions are estimated by replacing 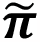 with *π*^(*b*)^ in the above steps.

### Trait goodness of fit

We quantify goodness of fit of our predictive trait model in two ways. First, we calculate Bayesian R-squared values for each functional trait considered in our analysis using a modified approach to that presented in Gelman et al., (2019). Bayesian R-squared values are calculated as 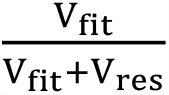 where V_fit_ and V_res_ indicate the fitted and residual variance, respectively (Gelman et al. 2019). Fitted variance is calculated as the variance among community-weighted mean prediction, 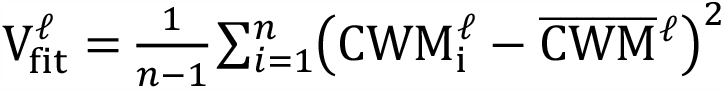, where CWM is calculated as described under Functional trait prediction and 𝓁 refers to a specific trait. Residual variance is calculated as the mean of the marginal variance of each trait across sites as calculated within the community weighted trait covariance matrix: 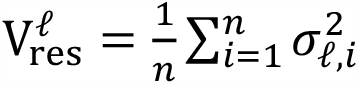. The Bayesian R-squared provides a measure of the variability in traits among sites explained by the model.

We further quantified the difference in observed versus predicted trait distributions across all sites applying a modified Kolmogorov-Smirnov test. This approach is novel in that it considers the full probability density function of the multivariate trait distribution rather than comparing observed versus predicted summary statistics (e.g., community weighted means, variances, coefficient of variation). We generate empirical cumulative density functions, *F*_𝓁_(*x*), for each modeled trait conditional on observed (***y***_*i*_) and predicted 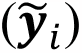 local communities. Specifically, we generate samples of species-level trait means with frequencies matching observed and predicted relative abundances of each species. For the predicted trait distribution, we apply the posterior predictive mean of the relative abundance of species within a local community sampled from a multinomial distribution with size equal to *N*_*i*_ and species probabilities equal to 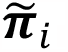. We then conduct Kolmogorov-Smirnov (KS) tests for each functional trait of interest across all sites with total observed abundance greater than 20 (*N*_*i*_ ≥ 20). A total of 3135 out of 3602 met this criterion (the consistency of KS tests is poor for low sample sizes). We applied a Bonferroni-adjusted test level of 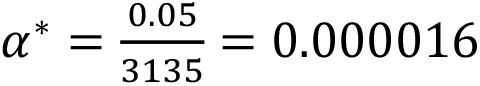 to control for multiple comparisons. The proportion of the 3135 sites for which there is evidence to reject the null hypothesis is reported for each trait of interest in Appendix S1: Table S2. Across all traits of interest, we find that the null hypothesis that the observed and predicted functional trait distributions are identical is supported at over 90 percent of sample sites.

### Species richness

To provide context for changes in functional trait distributions along modeled environmental gradients, we estimate species richness under specified environmental conditions. Specifically, we convert predicted species probabilities to presence/absence as: 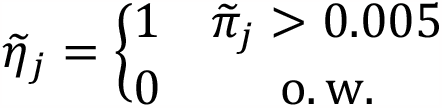. Species richness (*M*) is then estimated as, 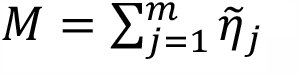.

## Results and discussion

### Model validation

We compared our model predictions to observed natural communities at two levels: 1) community composition, 2) functional trait distribution. In-sample and out-of-sample posterior predictive model checks revealed high correlation between predicted and observed relative abundances of species with species-level Bayesian R-squared values exceeding 0.6 for the majority of species (Itter et al. 2023). At the functional trait level, observed and predicted functional trait distributions were not found to differ significantly in over 90 % of study sites based on a modified Kolmogorov-Smirnov test (Appendix S1: Table S2). Clonality, SLA and leaf C:N were the best predicted traits, with the model capturing on average 28, 27 and 23 percent of their variation across over 3000 study sites, respectively (Appendix S1: Fig. S1). Further, community-level trait predictions (community weighted trait mean, CWM) correlated well with the observed CWM values for the best predicted continuous traits while the model poorly predicted variation in height (Appendix S1: Fig. S2).

### Community trait niche volume and species richness

Our analyses revealed that a reduction of community trait niche volume was generally associated with decreasing species richness in both low and high fertility habitats (Fig. 2, contour lines). This relationship can originate from random sampling (more species - greater trait niche; Cornwell et al. 2006), or be driven by ecological selection, caused for example by limiting similarity (Macarthur and Levins 1967). While our observational data does not allow disentangling the causes of this general relationship, an exception to this pattern points towards an ecological explanation: we found indication that high resource availability may allow a higher number of species to pack into a community without increasing its trait niche volume. Specifically, in the most fertile habitats with warmest summers, the maximum species richness did not coincide with the maximum trait niche volume, but richness peaked at low forest density while the community trait niche peaked at intermediate forest density (Fig. 2b). In the low-density forests, light and soil nutrients are abundantly available for understory vegetation, as the overstory tree layer is developing and not yet competing for resources as strongly as at intermediate and later successional stages. Nitrogen availability, in particular, has been shown to decrease as forests age (Merilä et al. 2002, Högberg et al. 2017). Previous research has shown that high nitrogen availability allows tighter packing of species with narrower nitrogen niches thereby enabling higher species richness (Heikkinen and Mäkipää 2010). Our analysis of multiple traits and their correlation suggests that also weaker trait correlation (Fig. 3j) and greater variation in foliar nutrients (C:N and P, Figs 3h,i) may permit different trait syndromes and thus higher species richness without increasing the total community trait niche volume. Similar strong niche packing was reported previously based on tree assemblages along a latitudinal gradient by Lamanna *et al*. (2014), who found that tropical tree assemblages hosted higher number of species within a smaller community trait niche than larger trait niches of temperate tree assemblages did. Stronger species packing inside a smaller total community trait niche suggests an increase in functional redundancy, *i*.*e*. presence of functionally similar species, which is known to enhance ecosystem stability (Walker 1995, Fonseca and Ganade 2001, Cadotte et al. 2011)

**Figure 2.**
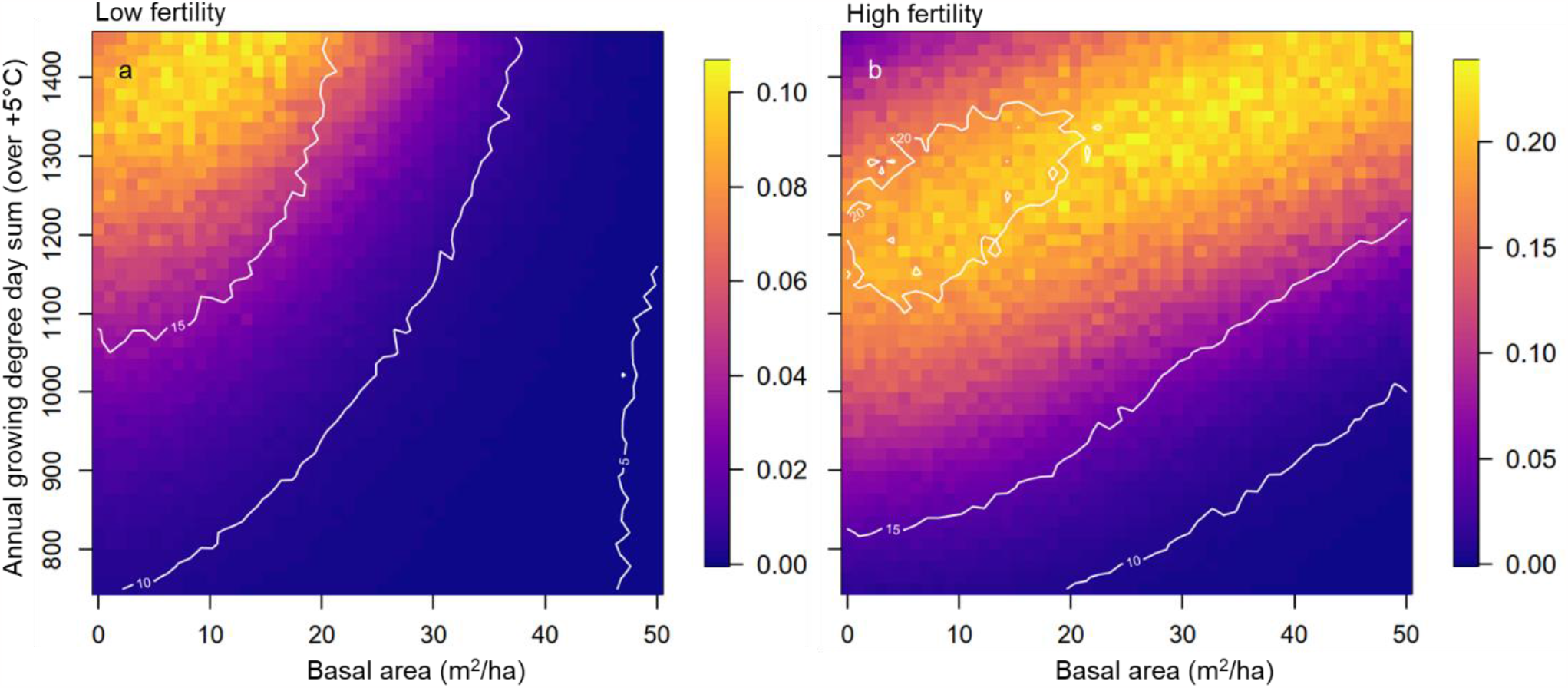
Four-dimensional community trait niche volume along environmental gradients. Relative change in community trait niche volume relative to baseline hypervolume, that assumes all species have equal probability of occurrence, along gradients of forest density (basal area of trees, x-axis) and growing season temperature (growing degree days, y-axis) in low (a) and high (b) fertility habitats. Values indicate proportion of the baseline volume (e.g., 0.1 = 10% of baseline volume). Note that colours refer to different values in the two panels. Contour lines represent species richness of predicted communities.

**Figure 3.**
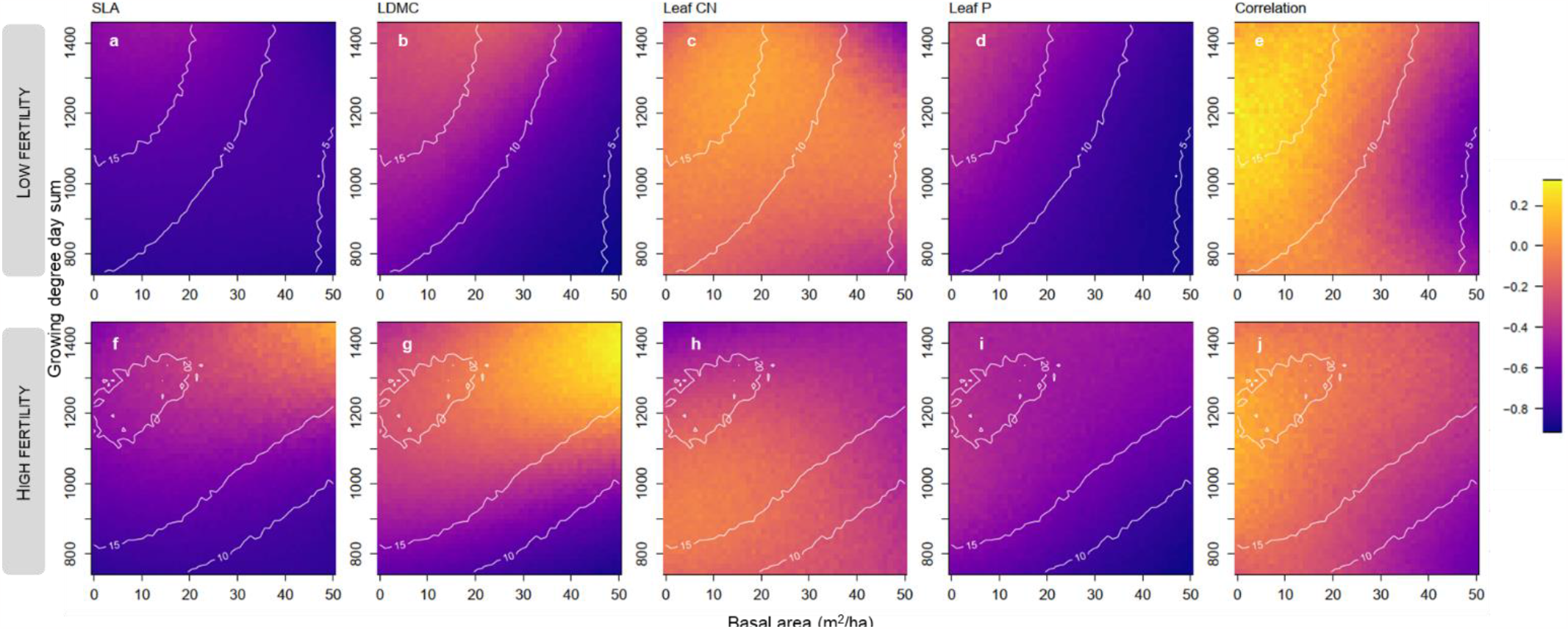
Functional trait variance and correlation among all traits along environmental gradients. Relative change in single trait variance and trait correlation along gradients of forest density (basal area, x-axis) and growing season temperature (growing degree days, y-axis) in low (top row) and high (bottom row) fertility habitats. Changes are calculated comparing predicted variance and correlation components to a baseline hypervolume that assumes all species have equal probability of occurrence. Positive values for single trait variances indicate an increase in variance relative to the baseline, while negative values indicate the opposite. For trait correlation, positive values indicate weaker correlation among traits in predicted communities compared to baseline community, while negative values indicate stronger trait correlation.

### Divergent selection

As succession proceeds and forest density increases, competition for nutrients increases and the resources for understory communities decline. Along the successional gradient, in warmer and more fertile habitats, we observed an expansion of the community trait niche (Fig. 2b). This was driven mainly by increasing variation in LDMC and to lesser degree by SLA (compare Fig. 3a-b to Fig. 3f-g). Simultaneously, we observed the LDMC distribution shifting from one dominant trait value to a more uniform distribution (Fig. 4j-k), corresponding to the pattern predicted to emerge from divergent trait-based selection (Vellend 2016). LDMC is related to resource use strategy and leaf lifespan describing species position on the resource use axis (Wilson et al. 1999). Thus divergent selection for LDMC, as succession proceeds, presumably reflects negative interactions among species, potentially intensifying competition for soil nutrients such as nitrogen, and favouring species with different resource-use strategies (Hyvönen et al. 2008, Högberg et al. 2017). This finding adds needed evidence to little studied divergent selection in real-world communities (Kingsolver and Pfennig 2007, Rolhauser and Pucheta 2017, Loranger et al. 2018).

**Figure 4.**
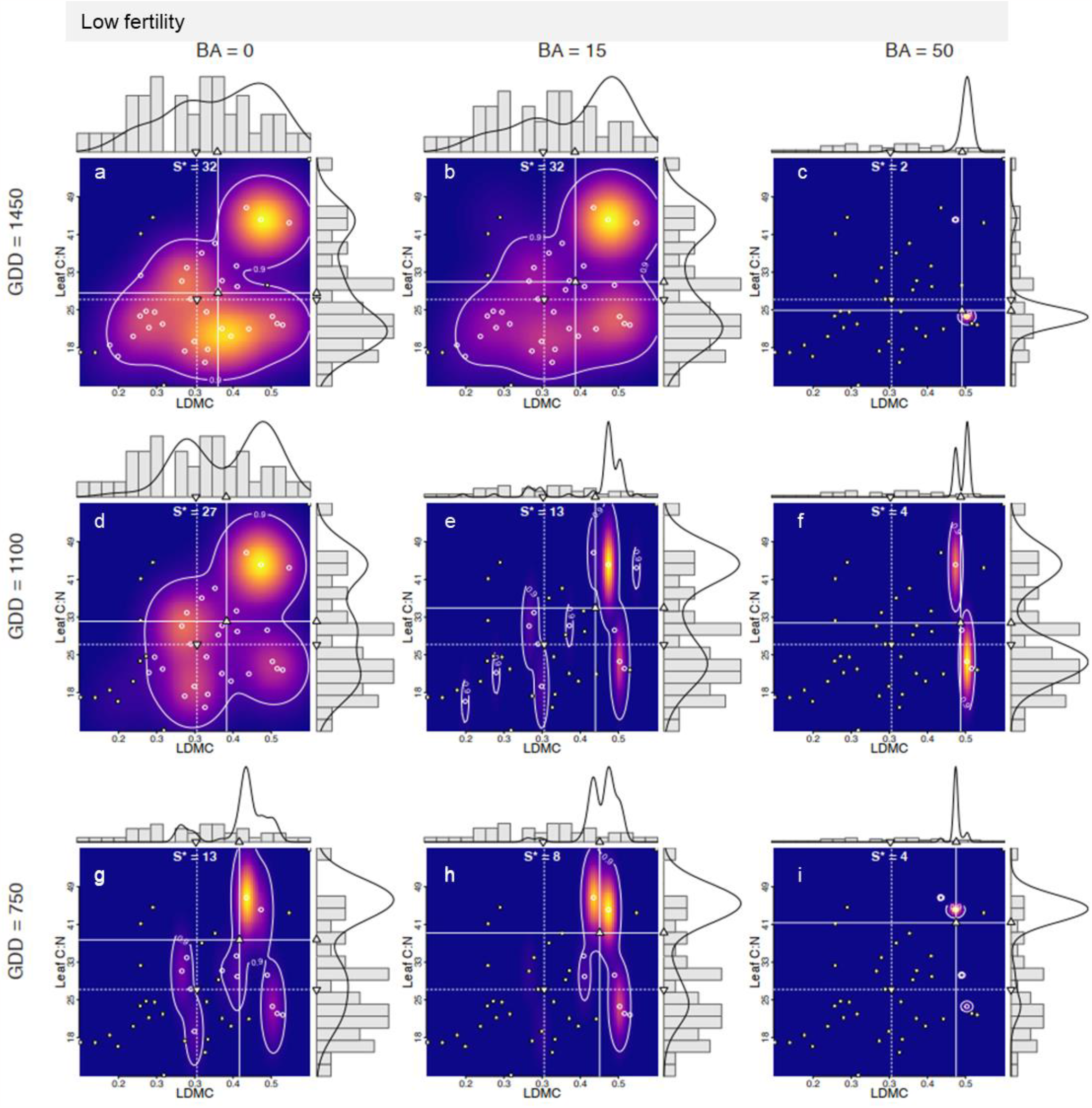
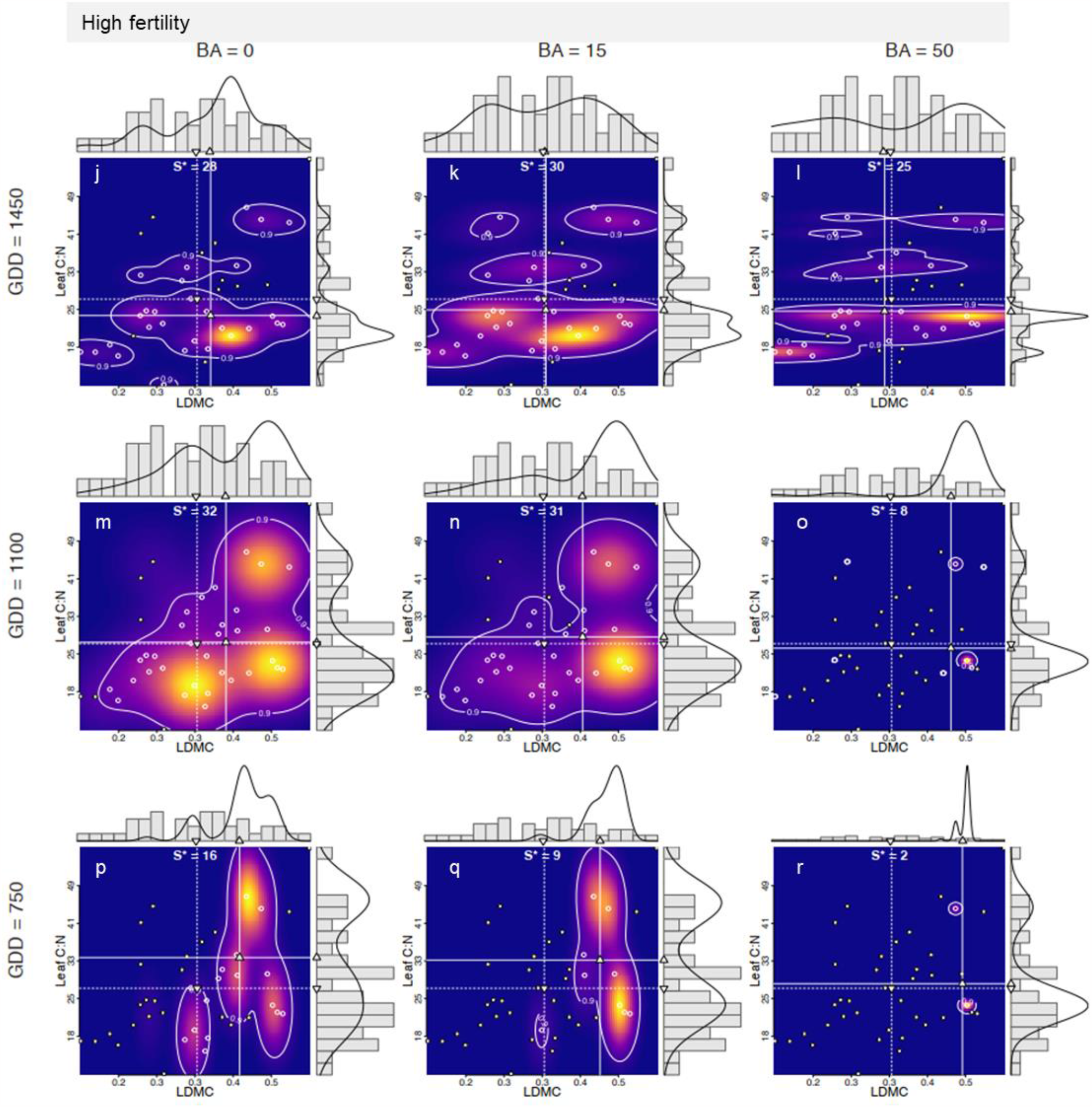
Empirical bivariate trait distributions along environmental gradients. Predicted empirical distribution of leaf dry matter content (LDMC, x-axis) and leaf C:N ratio (y-axis) under varying basal areas (columns) and growing degree days (rows) as predicted by the model in low (upper nine panels) and high fertility habitats (lower nine panels). In each panel, points indicate the 39 study species in the trait space (species identities in Appendix S1: Fig. S5) while the white line indicates the 95 % probability density of two traits. Species falling within the 95 % area are indicated with solid white symbols and their number with S* at the top. Modeled marginal community trait distributions are shown as solid black lines. Grey histograms present trait frequency distribution assuming all 39 species have equal probability of occurrence. The downward triangle in the margin and dotted lines denotes community-weighed mean (CWM) for the baseline community, while the predicted CWM is denoted by an upward triangle and dashed lines.

### Directional and stabilizing selection

In addition to divergent selection under certain conditions (warm, fertile sites), the predicted trait distributions revealed patterns resembling the hypothesized outcomes of directional and stabilizing selection (Fig. 1). For example, as an indication of directional selection, we observed an increase in the community weighted mean of leaf C:N when moving from warmer to colder growing seasons (Fig. 4). This pattern was most pronounced in the densest forest in low fertile habitats (Fig. 4c, f, i), suggesting that growing season length (or some correlated but unmeasured variable) acts as a selection agent favouring species with high nitrogen requirement (low leaf C:N) in warm conditions and species requiring less nitrogen in cold conditions. Beyond specific growing season length dependence, N availability may generally act as the proximate selection agent in boreal forests where decomposition rate (Kirschbaum 1995, 2006) and N mineralization (Liu et al. 2017) are strongly limited by temperature. Pure stabilizing selection, by definition, takes place when there is no change in the mean, but variance decreases, as we observe for LDMC in warm low fertile habitats (Fig. 4a-c) as succession proceeds (also observed for SLA, Appendix S1: Fig. S4a-c).

### Stabilizing selection due to trait correlation

Our results also indicated stronger correlation among traits along the successional gradient across habitat types (Fig. 3e, j). The increase in trait correlations in older (and, in fertile habitats, also colder) forests implies that the community trait niche is smaller than that defined only by variation in single traits, *i*.*e*., the dependence among traits shrinks the trait niche volume. Increasing trait dependence points to multiple concurrent assembly processes inducing trait trade-offs across the modelled gradients. Trade-offs limit the number of species with viable trait combinations especially in the densest and coldest forests. An earlier study reported increasing covariation of two traits, height and seed mass, along an aridity gradient (Dwyer and Laughlin 2017). In support of this, our results revealed changing dependence among traits along multiple interacting gradients, and add to emerging evidence of multiple simultaneous assembly processes along environmental gradients (Cornwell and Ackerly 2009, Spasojevic and Suding 2012). At the same time, our finding contradicts the common view that communities change from functionally and phylogenetically clustered to overdispersed as succession proceeds (Meiners et al. 2015, Cadotte and Tucker 2017). Overall, we found clear evidence that functional diversity of understory communities decreases along a successional gradient in this system characterized by a wide range of growing season temperatures and habitat fertility variation.

### Shifts in selection types along a successional gradient

We found evidence that multiple selection types modify community trait niches over the course of succession. First, we observed divergent selection on leaf C:N from the low-density (Fig. 4a) to mid-density forests (Fig. 4b) in warm low-fertile forests, and later stabilizing selection on the same trait, when moving to the oldest and densest forests (Fig. 4c). Here, the underlying mechanisms behind divergent selection is likely to be competition for nitrogen (Heikkinen and Mäkipää 2010), while stabilizing selection in the densest forest could be driven by competitive exclusion by the dominant species *Vaccinium myrtillus* (Gundale et al. 2012), although we cannot exclude other possible mechanisms (Table 1). To our knowledge, this is one of the first illustrations that at community-level, ecological selection on one trait may change over time, shedding light on one of the open questions regarding phenotypic selection in the natural communities (Kingsolver and Pfennig 2007).

### Advancing detection of assembly processes

Our approach improves identification of potential community assembly processes in three ways. First, because our framework builds on modelling species’ relative abundances within communities, it accounts for inherent trait correlations (or trait syndromes) within species and enables prediction of community trait composition under selection, which ultimately acts on individuals, but scales up to community-level trait patterns (Webb et al. 2010, Vellend 2010, Rolhauser and Pucheta 2017, Munoz et al. 2018). Second, our modelling approach allows for non-parametric trait-abundance distributions including skewed, peaky and flat distributions, thus more accurately reflecting real-world communities (Rolhauser and Pucheta 2017, Falster et al. 2021). Third, our model allows multimodality in trait distributions, which enables detection of divergent selection and shifts in modality, which so far have remained largely undetected (Loranger et al. 2018). Recent research has focused on improving detection of assembly processes by using multiple aspects of community trait distributions (e.g. skewness and kurtosis) as opposed to traditional mean-variance dominated methods (Laughlin et al. 2015, Gross et al. 2021) which implicitly assume Gaussian trait distributions. Our approach advances this field by demonstrating shifts in probabilistic community trait niches, and by linking them to ecological selection types, potential underlying low-level mechanisms, and species diversity patterns.

## Conclusions

We introduced a probabilistic model-based framework to build multidimensional community trait distributions and illustrated how they can be used to detect signs of ecological selection and to identify possible underlying mechanisms. By implementing this framework on a large observational dataset, we found evidence that directional, stabilizing and divergent selection concurrently shape boreal forest understory communities along environmental gradients. Our results also illustrated that the type of selection acting on a trait can change from divergent to stabilizing over time, as succession proceeds. Finding that multiple selection types are possible over time, brings novel insight to ecological theory, which often assumes that traits are either under stabilizing or directional selection (Kingsolver and Pfennig 2007, Vellend 2016). In these successional communities, interacting and alternating selection forces created multiple trait trade-offs selecting for species with specific trait combinations. As a result, we observed a reduction in the overall functional diversity and species richness of boreal forest understory communities along the successional gradient. All these findings remain to be validated with experimental data, which allows separation of selection, drift, and dispersal. We encourage researchers with suitable experimental data to test the robustness of this framework in detecting experimentally manipulated selection correctly. Taken together, our results illustrate how interacting selection processes acting on individuals representing multidimensional phenotypes leave distinct imprints on trait niches and the diversity of communities.

## Acknowledgements

We thank all people who participated in field surveys and managed the data at Natural Resources Institute Finland (Luke). This study was enabled by funding from Jane and Aatos Erkko Foundation to the Research Centre for Ecological Change (REC) and grants from Finnish Cultural Foundation and Oskar Öflund Foundation to EK, and Academy of Finland (grant 347188 to EK, and 317255 to JV). Natural Resources Institute Finland funded the work of TT, LH, MS and PM. The sampling in 2006 was co-funded by the BioSoil project carried out under the Forest Focus scheme [European Commission Regulation Nr. 2152/2003]. For part of the trait data, we acknowledge the TRY plant trait database which is hosted, developed and maintained at the Max Planck Institute for Biogeochemistry by J. Kattge and G. Bönisch.

## Author contributions

EK and MI developed the conceptual and theoretical framework and contributed equally. TT, LH, MS and PM were responsible for field sampling and data curation. MI and JV developed the model. MI performed data analyses and coding. EK wrote the first version of the manuscript, and all co-authors contributed to revisions.

